# Transcription through the eye of a needle: daily and annual cycles of gene expression variation in Douglas-fir needles

**DOI:** 10.1101/117374

**Authors:** Richard Cronn, Peter C. Dolan, Sanjuro Jogdeo, Jill L. Wegrzyn, David B. Neale, J. Bradley St. Clair, Dee R. Denver

## Abstract

**Background:** Perennial growth in plants is the product of interdependent cycles of daily and annual stimuli that induce cycles of growth and dormancy. In conifers, needles are the key perennial organ that integrates daily and seasonal signals from light, temperature, and water availability. To understand the relationship between seasonal rhythms and seasonal gene expression responses in conifers, we examined diurnal and circannual needle mRNA accumulation in Douglas-fir (*Pseudotsuga menziesii*) needles at diurnal and circannual scales. Using mRNA sequencing, we sampled 6.1×10^9^ microreads from 19 trees and constructed a *de novo* pan-transcriptome reference that includes 173,882 tree-derived transcripts. Using this reference, we mapped RNA-Seq reads from 179 samples that capture daily, seasonal, and annual variation.

**Results:** We identified 12,042 diurnally-cyclic transcripts, 9,299 of which showed homology to annotated genes from other plant genomes, including angiosperm core clock genes. Annual analysis revealed 21,225 an-nually-cyclic transcripts, 17,335 of which showed homology to annotated genes from other plant genomes. The timing of maximum gene expression is associated with light quality at diurnal and photoperiod at annual scales, with two-thirds of transcripts reaching maximum expression +/− 2 hours from sunrise and sunset, and half reaching maximum expression +/− 20 days from winter and summer solstices. Comparison to published microarray-based gene expression studies in spruce (*Picea*) show that the rank order of expression for 760 putatively orthologous genes was significantly preserved, highlighting the generality of our findings.

**Conclusions:** This finding highlights the extensive annual and seasonal transcriptome variability demonstrated in conifer needles. At these temporal scales, 29% of expressed transcripts showed a significant diurnal rhythm, and 58.7% showed a significant circannual rhythm. Remarkably, thousands of genes reach their annual peak activity during winter dormancy, a time of metabolic stasis. Photoperiod appears to be a dominant driver of annual transcription patterns in Douglas-fir, and these results may be general for predicting rhythmic transcription patterns in emerging gymnosperm models.

## BACKGROUND

The sensing of daily, seasonal, and annual environmental variation in land plants is accomplished using a diverse array of organs, transcriptional regulators that drive oscillatory functions, and pathways that refine, optimize, and entrain rhythms to rhythmic environmental stimuli. Circadian patterns are ubiquitous in photosynthetic [1][2][3][4] and non-photosynthetic [5][6] organisms, and they are essential for coordinating external signals for optimally timing transcriptional activity to match the demands of growth and phenology with resource availability. The genetic basis for diurnal responses in model plants have been intensively studied using mutant screens [4][7][8] and global transcription [7], and these studies have identified components of the central oscillator (“core clock”) and genes that are targets of rhythmic activation and repression. Variability in core clock genes alters the timing of the clock and clock-dependent pathways, so subtle changes in genes controlling clock rhythm may contribute to local adaptation on a short time scale, and evolutionary divergence on a longer time scale [1][9].

In temperate zone trees, circadian rhythms are superimposed on longer annual cycles involving transitions between active growth -- the time when light energy is captured and converted into growth -- and dormancy, the time when growth potential is arrested to protect cells from seasonal stresses of cold temperatures, freezing and desiccation [10][11][12]. Accurately timed transitions between growth and dormancy are essential for adapting to variable environments [13] and they depend on: (1) reliable environmental cues that forecast future change; (2) diverse sensory organs; and (3) competing biochemical networks that integrate sensory signals and shift responses from one ‘module’ (growth; dormancy; senescence) to the next. Photoperiod and light quality are known to be important cues for initiating seasonal growth rhythms and establishing the onset of dormancy for many trees [11][12][14][15], so pathways involved in light capture and photoperception are expected to show annual rhythmic variation. As dormancy is established, trees sense and respond to cold temperatures (chilling units) by increasing cold hardiness and increasing resistance to desiccation. After chilling thresholds are met, trees respond to warming temperatures (forcing units) by releasing tissues from dormancy, remaining in a ‘stand-by’ state until requirements for initiating growth (heat, water availability, light quality [10][14][16]) are met. Genes implicated in dormancy and resumption of spring growth should also show annual cyclic variation, and these include components of the circadian clock and photoperiod-responsive genes [10] [11][14][15], and pathways involved in temperature and water perception [15][17][18], hormone regulation and cell growth [10][19], and glucan hydrolysis [19].

The organs responsible for sensing and integrating external stimuli -- leaves, shoots, and roots -- are common to perennial plants, but signal perception and integration during the dormant season is likely accomplished by different means in conifer trees with perennial leaves (“needles”), versus trees with deciduous leaves. Perennial needles confer advantages by preserving annual investments in carbon fixation, conducting photosynthesis year round [20][21], offering alternative mechanisms for preventing winter embolisms [22], and providing an environmentally-responsive sensor that adds to bud- and stem-associated signals during dormancy. Perennial needles also come with fitness trade-offs because they can be damaged by cold during entry into dormancy and during de-acclimation in spring [21] [23], and by photosystem excitation that can lead to the formation of reactive oxygen species during dormancy [21] [24]. Given the complexity involved in orchestrating growth and stasis over annual cycles, annual gene expression variation in conifer needles should show high complexity, especially when compared with the annual leaves characteristic of model trees (*Populus* L.) and herbs (*Arabidopsis* Heynh. in Holl & Heynh).

To gain an understanding of the complexity of circadian and circannual rhythms of gene expression in conifers, we examined cyclic gene expression in needles from Douglas-fir (*Pseudotsuga menziesii* (Mirb.) Franco) at daily and annual scales. Douglas-fir is related to model conifers like Norway spruce (*Picea abies* L.) and Loblolly pine (*Pinus taeda* L.) through divergence in the Early Cretaceous ~130 MYA [25], and to angiosperms like *Populus* and *Arabidopsis* through a more ancient divergence ~300 MYA. Douglas-fir has an expansive native range in North America [26], and it is noteworthy among conifers for its significant population variation in needle cold hardiness, phenology, and growth traits [27] [28]. Some of the strongest associations between provenance source and quantitative traits in Douglas-fir are exhibited by needle traits related to annual cycles of freeze-avoidance, such as spring and fall needle cold hardiness, and rhythmic cues that define the onset of winter like the first winter freeze and variability in the frost free period [23][29][30].

In this study, we use next-generation mRNA sequencing to produce individual *de novo* needle transcriptomes from 19 Douglas-fir trees, and use the resulting “pan-transcriptome” as a reference for mapping RNA-seq reads from experiments evaluating diurnal variation over two daily cycles and circannual variation over one annual cycle. We specifically searched for transcripts exhibiting rhythmic expression [31][32] to define the timing of maximum expression (“phase”) and amplitude. Our results provide the first characterization of year-round transcriptome-wide activity in leaves from a perennial plant, and they identify a core set of transcripts that show evidence for significant cycling on seasonal and annual scales. Our results show congruence with microarray-based studies of seasonal gene expression in Sitka spruce [33], and they underscore the potential for using gene expression information from Douglas-fir to predict cyclic transcriptome responses, and gene expression patterns responding to different environmental cues in temperate-zone conifers.

## METHODS

### Plant materials and sample information

Trees used for annual analysis are from a larger reciprocal translocation study [18] [34] that includes multiple sources of *Pseudotsuga menziesii* var. *menziesii* from the Pacific Northwest of North America. Trees were chosen to maximize differences in source climatic and phenology [28]; included are families from regions that derive from cold/wet sources (47.3 ° N, −121.6 ° W, 950 m elevation), cool/wet sources (47.2° N, −123.9 ° W, 111 m elevation), and warm/dry sources (43.3 ° N, −123.1 ° W, 429 m elevation; supplemental Notes S1). Two-year old trees were planted in a warm/dry region (Central Point, Oregon, USA; 42.3° N, −122.9° W, 390 m elevation) in November, 2009, and sampling of the 2010 cohort of needles was initiated on October 27, 2010 at ~ three week intervals until November, 2011 (16 samples points; Fig. 1). Needles were collected between 11am and 1 pm (ZT = 05:00 to 07:00) from 13 individual trees. Environmental data (sunrise; sunset; day length; cumulative weekly precipitation; minimum and maximum daily temperature) was collected over the duration of this experiment (File S1; Fig. 1). Sampling intervals and RNA-seq sample sizes for the annual study are summarized in Notes S1. Trees used for diurnal analysis derive from the warm/dry region (43.3 ° N, −123.1 ° W, 429 m elevation) and were grown in Corvallis, Oregon, USA (42.3 ° N, −122.9 ° W, 74 m elevation; supplemental notes S1). Needles were collected from 6 individual three-year old trees three half-sib families, two sibs per family). Needles were collected at 4 hour intervals, starting at 2 AM, for a total of 48 hours in early fall (September 7, 8). For this experiment, sunrise (ZT0) occurred at 06:44 AM, and sunset occurred at 19:34 PM, giving a 12:50 photoperiod. Sampling intervals and RNA-seq sample sizes for the diurnal study are summarized in Notes S1.

**Figure 1.**
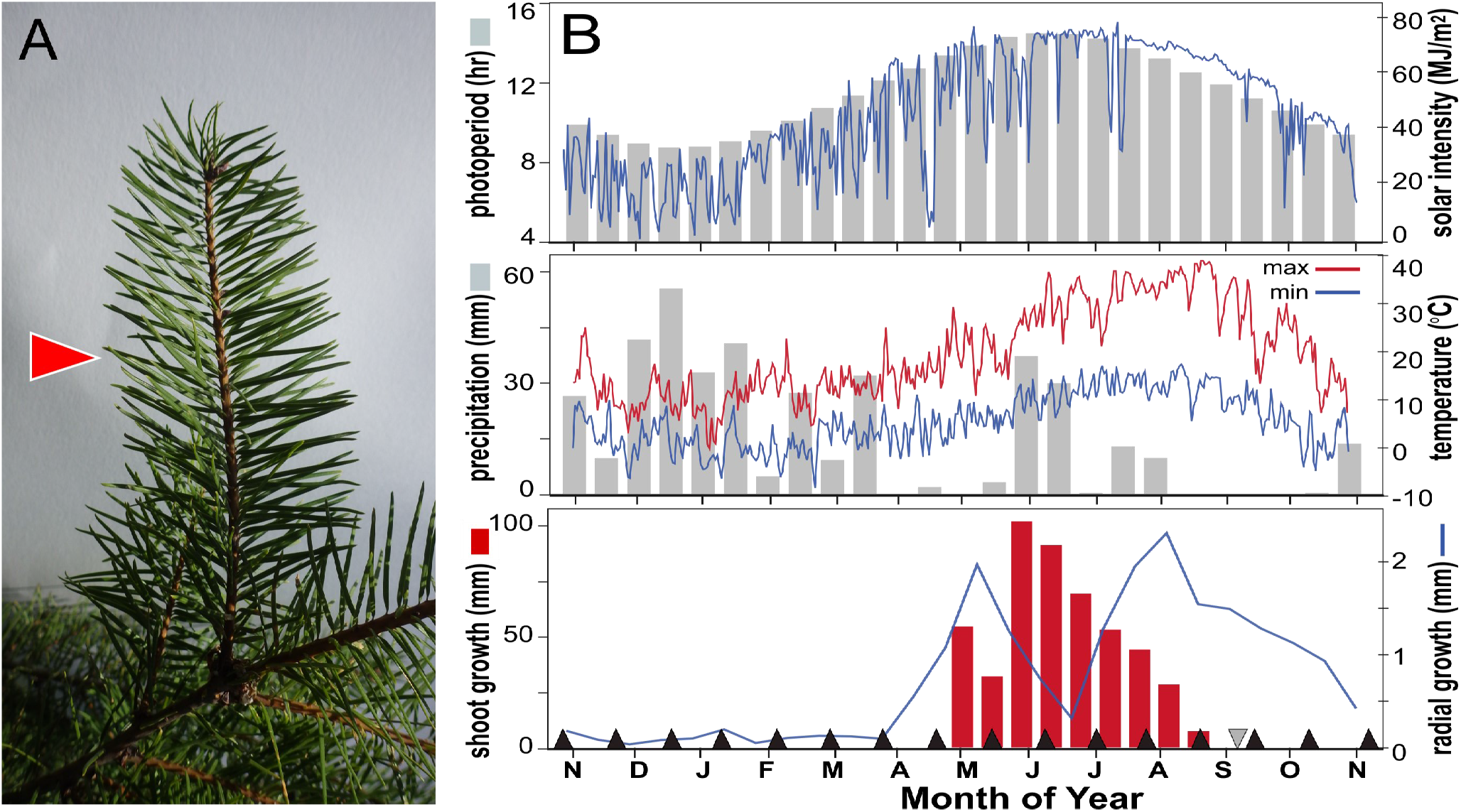
Inputs for the diurnal and annual needle transcriptome study. (A) Douglas-fir branches, showing the location of sampled needles (red arrow). (B) Annual environmental conditions and growth rhythm for trees used in this experiment. Shown are: upper panel, light conditions with photoperiod (bar chart) and solar incidence (blue line); middle panel, weather conditions with precipitation (bar chart) and temperature (high = red line; low = blue line); lower panel, growth rhythm with terminal shoot growth (bar chart) and radial growth (blue line). Annual needle sample times are identified in the lower panel using black triangles. The date of the diurnal study is identified as a grey triangle.

### Needle collection, RNA isolation and RNA sequencing

Needles (8 – 12; Fig. 1) were collected from branches representing four cardinal directions of a single tree, flash-frozen in liquid N_2_, and ground at dry ice temperatures (FastPrep-24 mill; MP Biomedical, Solon, OH, USA). Ice-cold extraction buffer (1.5 mL of 3M LiCl/8M urea; 1% PVP K −60; 0.1M DTT [35]) was added to ground tissue, homogenized, then centrifuged at 200g × 10 min., 4 ° C. The supernatant was incubated overnight at 4° C, and crude RNA was pelleted (20,000g × 30 min., 4°C) and cleaned using the ZR Plant RNA MiniPrep kit (Zymo Research, Irvine, CA, USA) and DNase treatment (Turbo DNase; New England Biolabs, Ipswich, MA, USA). RNA concentration was estimated using a Qubit fluorometer (Invitrogen, Carlsbad, CA, USA), and RNA quality was checked using an Agilent BioAnalyzer (Agilent, Santa Clara, CA, USA).

Indexed Illumina RNA-seq libraries used 2 μg total RNA and TruSeq chemistry (Illumina Inc., San Diego, CA, USA), modified for strand-specific sampling [36]. In this protocol, first-strand synthesis products were desalted to remove unincorporated dNTPs (Sephadex G-25; Sigma-Aldrich, St. Louis, MO, USA), and reconstituted in dNTP-free second-strand synthesis buffer with second strand enzyme mix (New England Biolabs) and a dUTP/dNTP mixture (Thermo-Scientific, Waltham, MA, USA) to incorporate dUTP into the second strand. All other steps follow Illumina protocols, except that uracil-containing strands were degraded using a uracil-specific excision reagent mixture (37°C for 15 minutes; New England Biolabs) prior to PCR. Amplified libraries were quantified and pooled at equimolar 6-plex representation at 10 nM. Sequencing was performed at Oregon State University’s Center for Genome Research and Biocomputing (Corvallis, OR, USA) using a HiSeq 2000 (Illumina Inc.) with version 3.0 chemistry and demultiplexing performed using Casava v1.8 (Illumina Inc.). The experiment included 179 libraries sequenced on 34 lanes from 10 different flow cells, and it is recorded under NCBI Umbrella BioProject PRJNA362352; see Notes S1).

### Transcriptome assemblies and annotation

Microreads from individual trees were combined over all time periods to create 19 single-tree source files. Reads were quality trimmed using Trimmomatic v.0.30 [37] (using options `-phred33 LEADING:20 TRAILING:20 SLIDINGWIN-DOW:5:20`), and 19 individual transcriptome assemblies were de novo assembled using *Trinity* v.r2013_08_14 [38] using a minimum size of 200 bp. To create a pan-transcriptome reference, the longest transcripts from each component in single-tree de novo assemblies were identified and combined into a single file, and transcripts smaller than 300 bp were removed. This combined file was sorted by sequence length and then clustered using USEARCH v.7.0.1001 [39]. The `usearch64 - cluster_smallmem` command was used with a sequence identity threshold of 90% (-id 0.9 flag) and with the -`strand both` flag to combine forward and reverse transcripts into the same clusters. A table of the number of input bases from each library, transcripts assembled for each individual, and the combined clustered reference assembly is provided in Notes S1. Individual assemblies and the pan-transcriptome reference are available for download at the TreeGenes Forest Tree Genome database web site under the link “*Pseudotsuga menziesii* Transcriptome” [40].

To annotate plant-derived transcripts, we used BLASTX and TBLASTX [41] to identify transcript matches from the NCBI NR database (minimum identity; expect < 1e^−10^), and BLASTX to identify matches to the Mercator plant metabolic function database [42][43][44]. We used LASTZ [45] to identify chloroplast and mitochondrial transcripts using the *Pseudotsuga sinensis* chloroplast (NC_016064.1) and Loblolly Pine draft mitochondrial [46] genomes as references. We used BLAT [47] to search for conservation between Douglas-fir transcripts and conifer genome references, the Loblolly pine reference genome [48](Pinus *taeda* version 1.0), and a pre-publication draft for the Douglas-fir genome [49](version 0.5). LASTZ and BLAT searches used a match criterion of ≥80% identity with contiguous hits ≥ 50 bp. Annotations are available at at the TreeGenes Forest Tree Genome database web site under the link “*Pseudotsuga menziesii* Transcriptome” [40].

### Detecting diurnal and annual cyclic transcriptome variation

RNA-seq reads from individual tree samples were aligned using BowTie 2.2.3 [50] with the following call: `bowtie2 --end-to-end -D 15 -R 2 -L 22 -i S,1,1.15`. This allowed 15 consecutive seed extension attempts before the aligner moved on (-D 15), a maximum of 2 attempts to re-seed reads with repetitive seeds (-R 2), and a 22 bp seed (-L 22) with zero allowed mismatches in this seed. The function to determine the interval between seed substrings during multi-seed alignment was set to `f(x)=1+1.15* qrt(x)`, where x is read length (`−i S,1,1.15`); based on 101 bp read lengths, this resulted in an interval of 13 bp. For this experiment, transcripts showing a median ≤ 5 counts were considered background and were excluded from subsequent analyses.

For transcripts exceeding the detection threshold, the 72 diurnal samples (six individuals; 12 time points) were collected into one table and transcript counts were normalized using DESeq [51][52] [53]. Annual samples were similarly tabulated, median filtered, and DESeq normalized. After normalization, we computed family means by averaging reads from half-siblings (trees 44 and 90 = family A; trees 43, 46, and 49 = family B). For the annual study, counts were linearly interpolated to emulate equally -spaced sample intervals. The two processed count tables (diurnal, annual) were passed through JTK-cycle [31] to identify statistically significant rhythmic transcription (*p*≤0.05), after application of a false-discovery rate correction of *q*≤0.01 [54]. We used JTK-cycle to identify the phase (time point at which the underlying curve reaches its maximum value) for each transcript, with phases measured in hours after 12:00 AM for the diurnal study, or Julian days (days after January 1) for the annual study. Summaries of cyclic properties (phase; amplitude; period) are provided in supplementary file S2.

### Defining relationship between transcriptional phases and solar and weather factors, and enrichment/depletion tests by season

To evaluate the relationship between the timing of maximum annual gene expression (“phase”) and environmental variables, annual transcriptional phases were sorted into two week bins, starting on the first sample date (27-October-2010), and continuing until the last sample date. Counts of transcripts reaching maximum expression within two week bins were tallied, and counts were compared to environmental variable summaries for the same intervals; mean photoperiod (hours), the sum of precipitation (mm), and the mean low and high temperatures (°C). The number of genes reaching phase per two-week bin was modelled as a function of solar and environmental factors using linear and polynomial fits (degree = 2).

To evaluate Mercator metabolic terms for enrichment or depletion, we binned significantly cyclic transcripts (e.g., *q* ≤ 0.01 from JTK-cycle) from diurnal and annual experiments into four temporal “phase bins” of equal time duration. For the diurnal experiment, phase bins were six hours in length, with the first bin approximately centered on “sunrise” (4:01am – 10:00am), and subsequent bins defined as “midday” (10:01am – 16:00pm), “sunset” (16:01pm – 22:00pm), and “night” (22:01pm – 04:00am). Annual bins were 91 or 92 days in length, with the first bin approximately centered on the winter solstice or “short photoperiod” (5-Nov to 4-Feb), and subsequent bins defined as “spring photoperiod” (5-Feb to 5-May), “long photoperiod” (6-May to 5-Aug), and “fall photoperiod” (6-Aug to 4-Nov). Enrichment tests for Mercator pathway terms were performed using term lists for transcripts identified as significantly cyclic and identified to a function (all Mercator bins except 35.2, which is “not assigned.unknown undefined”). This resulted in lists of 7,415 annotated transcripts for the diurnal experiment (sunrise bin = 1,916; midday = 1333; sunset = 2,685; night = 1,481), and 13,514 annotated transcripts for the annual experiment (short photoperiod bin = 5,796; spring photoperiod = 1,466; long photoperiod = 5,073; fall photoperiod = 1,179). Enrichment/ depletion tests were performed using a one-tailed Fisher’s exact test and the program `Mefisto` [55]; adjustments for two-tailed tests were made by multiplying *P*-values by 2, as recommended by Rivals et al. [56], and a false-discovery rate correction of 1% (*q* < 0.01; [54]) was applied using the `p.adjust` function in the R library `stats`.

### Comparing annual transcriptome expression variation to other conifers

We compared our annual RNA-seq expression data from a subset of dates (17-August to 7-October) with a previously published microarray-based study of needle gene expression in Sitka spruce (*Picea sitchensis*) conducted over a similar time and season interval in British Columbia, Canada (30-August to 18-October)[33]. Putative orthologs of Douglas-fir and Sitka spruce transcripts were identified using reciprocal best BLASTN matches between the Douglas-fir pan-transcriptome reference and 18,237 clone sequences used in the spruce 21.8k microarray [33] We evaluated the conservation of expression patterns between studies by computing the ratio of fall:summer gene expression using transcripts showing significant expression differences (*q*<0.05) in microarray data [33] :

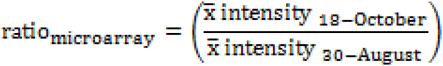

versus transcripts showing significant circannual rhythm (*q*≤0.01. from JTK-cycle) in RNA-seq data:

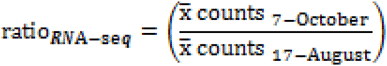

For the 760 transcripts meeting these criteria, expression ratios were ranked from high-to-low and ranks were compared by Kendall’s tau(τ) with the `cor.test` function in R. In this comparison, τ is bounded by +1 and −1, with the bounds representing perfect preservation of ranked gene expression ratios in the same (+1) or opposite (−1) direction, and 0 representing random ordering of gene expression ratios between experiments.

## RESULTS

### Defining the needle ‘pan-transcriptome’ of Douglas-fir

In this study, needle tissue was sampled by mRNA-Seq to evaluate diurnal and circannual variation in global transcription (Fig 1A, Fig 1B; supplemental notes S1). Needles were sampled for RNA at different time intervals to evaluate two transcriptome responses: (1) diurnal responses, using a sampling interval of four hours across two days (12 time points); and (2) circannual responses, using a sampling interval of approximately 3 - 4 weeks across a complete year (16 time points). This sampling scheme resulted in a data set that included 19 trees and 179 individual RNA-seq libraries to evaluate different aspects of temporal needle gene expression.

Individual tree mRNA-seq libraries from needles yielded 94.0 - 573.8 million reads, and individual *de novo* assemblies using *Trinity* produced 47,976 - 126,355 components 200 bp or larger (Table S2). The number of assembled *Trinity* transcriptome sequences and cumulative sequence length were positively and significantly correlated with the number of input reads (*r*^2^ ≥ 0.92; Notes S1). Across all assemblies, the majority of *Trinity* components (85.7%) showed a single subsequence, while the maximum number of sequences per component in any library was 227. A ‘pan-transcriptome’ reference was created using the longest sequence from each component in single-tree *de novo* assemblies, followed by clustering to reduce allelic and redundant sequences to one representative sequence. This step reduced the pool of transcripts from 1.66 million sequences from 19 individual-tree assemblies, to a pan-transcriptome with 199,471 sequences.

Multiple sources of evidence were used to characterize plant-derived transcripts for homologies and putative functions (Notes S1). BLASTX searches of the plant-specific Mercator plant metabolic database [42][43] identified homologies for 46,436 transcripts, while BLASTX and TBLASTX searches of NCBI NR database identified tentative identities for 54,384 transcripts. Homology searches against draft conifer genomes identified 102,714 homologs from the Loblolly pine v. 1.0 reference genome, and 167,821 homologs to the Douglas-fir v. 0.5 draft assembly. In total, these searches identified 173,882 transcripts (159.2 Mbp) as derived from Douglas-fir, with 143 originating from the chloroplast genome, 196 from the mitochondrial genome, and 173,544 originating from the nuclear genome. Of the remaining 25,589 transcripts, 19,000 were positively identified by BLAST as derived from foliar metaflora/metafauna present on Douglas-fir needles, or contaminants from the sampling/library construction process. A final list of 6,589 transcripts could not be identified using BLAST searches or searches of either draft gymnosperm genome assembly; due to their uncertain origin, these transcripts were omitted from subsequent analyses.

### Diurnal transcriptome variation in Douglas-fir tracks daily light/dark transitions

Experiments to detect diurnal cyclic transcriptome variation included six individuals (two sibs per family; three families) collected over twelve 4-hour intervals (Notes S1). After mapping reads from individual diurnal libraries to the Douglas-fir pan-transcriptome reference, 41,382 transcripts met our threshold for analysis of diurnal cycling (median mapped reads > 5; Table 1, supplemental file S2). Following DESeq count normalization, JTK-cycle identified 15,487 transcripts as showing significant cyclic diurnal expression at a false-discovery rate of 5% (*q* ≤ 0.05). The nonparametric test used in JTK-cycle can identify transcripts as significantly cyclic even if they show minute amplitudes, such as those that might result from circadian fluctuations in total RNA levels [57][58]. Since the magnitude of daily RNA fluctuation is unknown for conifer needles, we adopted a more stringent false-discovery rate of 1% (*q* ≤ 0.01) to identify ‘high-confidence” cyclic patterns; this reduced the number of diurnal transcripts to 12,042.

**Table 1.** Numbers of transcripts from Douglas-fir showing evidence of diurnal and annual cycling.

| TRANSCRIPTS |  |
| --- | --- |
| Total assembled | 199,623 |
| Total in genome | 173,882 |
| DIURNAL |  |
| Expressed (median > 5 counts) | 41,382 |
| Cyclic ($q \leq 0.05$) | 15,487 |
| Cyclic high-confidence ($q \leq 0.01$) | 12,042 |
| ANNUAL |  |
| Expressed (median > 5 counts) | 36,145 |
| Cyclic ($q \leq 0.05$) | 24,688 |
| Cyclic high- confidence ($q \leq 0.01$) | 21,225 |

The distribution of expression phase times (Fig. 2A) across all high-confidence diurnal transcripts shows a pronounced bimodal pattern, with the highest proportion of genes reaching maximum transcript accumulation near sunrise (ZT0 = clock hour 06:44 AM) and before sunset (ZT = 12:50, or clock hour 19:34 PM). This pattern has been shown in diverse plants and animals [5][7], and is explained as “expression anticipation” for the transition from dark to light in the morning, and light to dark in the evening. Our high-confidence diurnal transcripts include homologues to many of the known core clock genes from angiosperm models [4][7](Fig. 3; Table 2). We compared the timing of maximum expression for representative clock and seasonal genes in Douglas-fir to values reported for *Arabidopsis* [7] in controlled chamber experiments. We found a high degree of concordance in the phase for many of these genes (Table 2), which is striking given the different methods used and organismal divergence involved in this comparison. Transcripts encoding homologs of *circadian clock-associatedl (CCA1), cryptochrome1 (CRY1), constitutive photomorphogenic 1 (COP1), vernalization insensitive 3 (VIN3), reveille 1 (RVE1), gigantea (GI), timing of CAB expression 1 (TOC1), lux-arrhythmo (LUX)* and *early flowering 4 (ELF4.3)* all reached maximum expression within 3 hours of the reported maximal expression of *Arabidopsis* (Table 2). A smaller number of genes - specifically, *flowering locus T (FT), late elongated hypocotyl (LHY)*, and *Zeitlupe (ZTL)* - showed pronounced differences in phase from *Arabidopsis*. These differences may be due to incorrect assessments of orthology in gene families, but in the case of *FT*, it is likely due to divergent gene functions in gymnosperms and angiosperms, as has been previously suggested [14][59]. A noteworthy finding is that a large proportion of high-confidence diurnal cycling transcripts (38.4%; *N* = 4,627) have no homology to known proteins in Mercator and are annotated as “not assigned.unknown” (Fig. 2A); the functions of these genes cannot be assessed by sequence similarity-based inferences. This finding suggests that gymnosperms could possess novel clock-dependent components (genes or non-coding RNAs) that lack clear homologs in angiosperm model plants.

**Figure 2.**
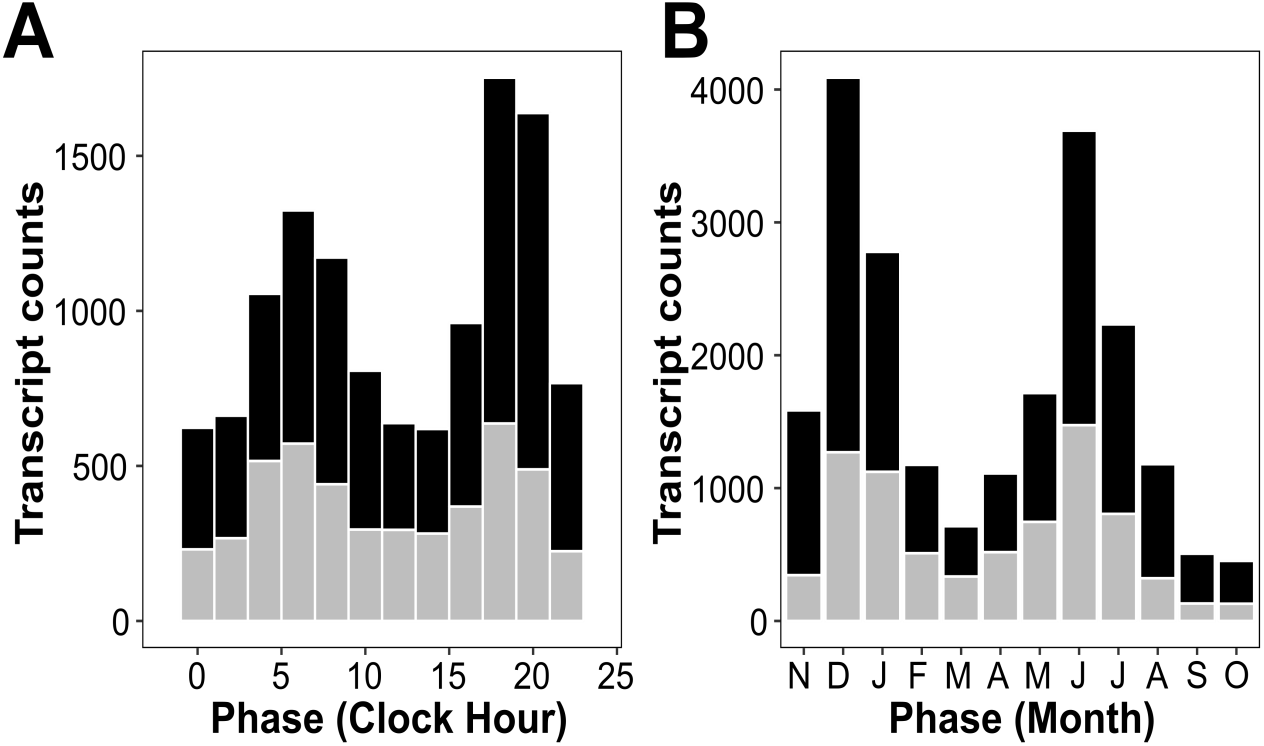
Frequency histograms of estimated phase times for transcripts showing cyclic expression patterns. (A) Histogram of phase for 12,042 transcripts showing significant diurnal cycling. Shown are counts per two hour interval, starting at clock hour 12:00 AM. (B) Histogram of phase for 21,225 transcripts showing significant annual cycling. Shown are transcript counts per one month interval, starting at November 1. In both plots, counts of transcripts with Mercator definitions are shown with black fill, while transcripts lacking Mercator definitions are shown with grey fill.

**Figure 3.**
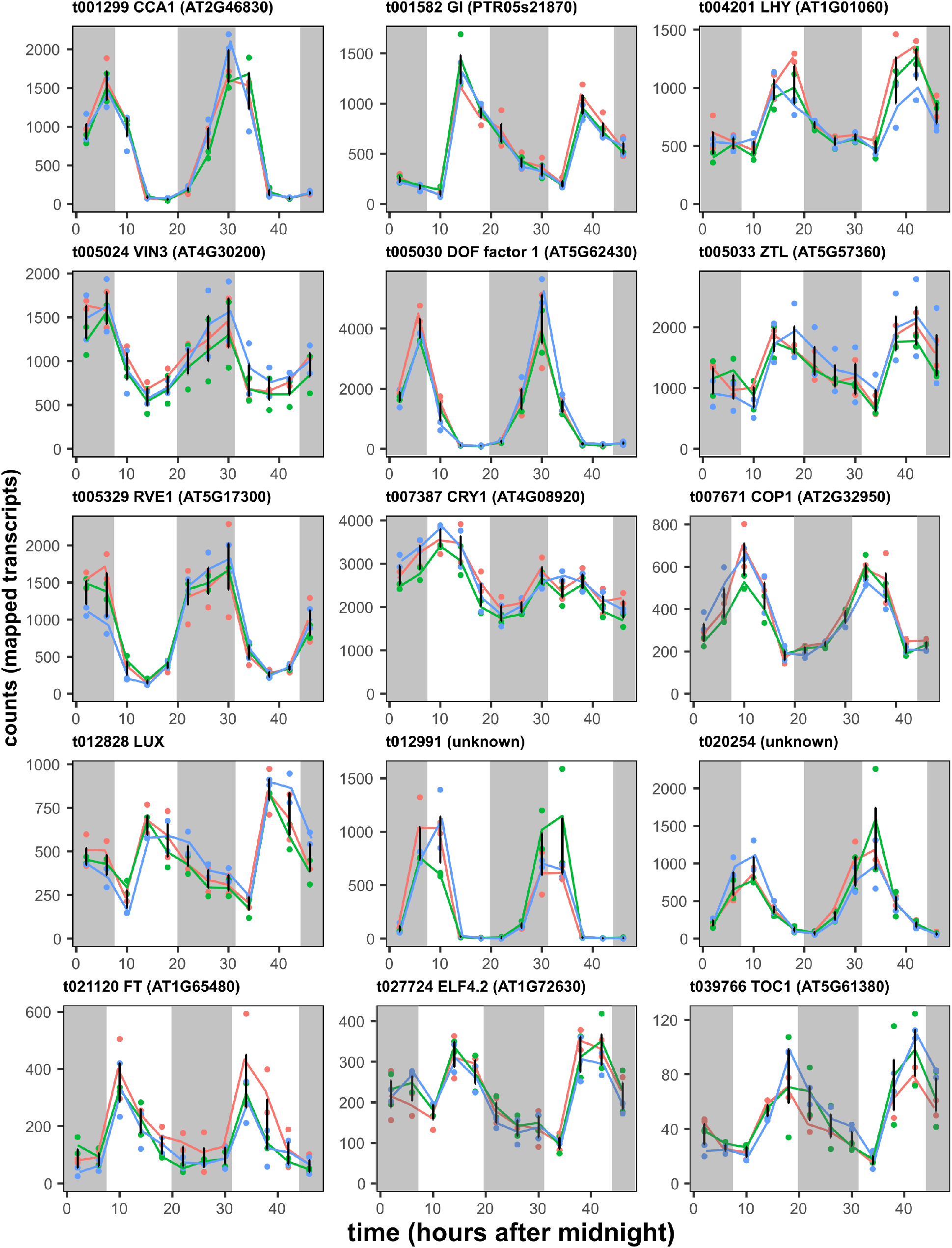
Example expression profiles for fifteen transcripts exhibiting significant cyclic diurnal variation, ordered by hourly phase. Gene expression (mapped counts per transcript) was estimated by RNA-seq from needle samples collected from three families of trees at four hour intervals for two days. For each gene, the Douglas-fir transcript name, putative gene name, and best BLAST match to *Arabidopsis* (prefix ‘at’) are provided. Lines connect the mean gene expression (DESeq-normalized counts per transcript) for each family, and error bars represent the SD for all replicates. Absence of shading indicates daylight hours, while shading indicates night hours.

**Table 2.** Diurnal and annual transcriptional phases for core clock genes in Douglas -fir needles, as compared to the angiosperm model *Arabidopsis*. Phases for Douglas-fir were computed using JTK_CYCLE and either the diurnal or annual data sets.

| Gene Description | Transcript or Locus |  | Diurnal Phase (clock time)<sup>2</sup> |  |  | Annual Phase<sup>2</sup> |  |
| --- | --- | --- | --- | --- | --- | --- | --- |
|  | Douglas-fir | *Arabidopsis* | Douglas-fir | *Arabidopsis* | Difference (hr) | Experiment Day | Month |
| *RVE1* (myb-like transcription factor) | t005329 | AT5G37260 | 0:00 | 3:00 | 3 | 97 | Feb |
| *VEL1* (vernalization insensitive-3-like) | t005024 | AT5G57380 | 4:00 | 3:00 | 1 | 255 | Jul |
| *CCA1* (circadian-clock associated) | t001299 | AT1G01060 | 4:00 | 6:00 | 2 | *n.s.*<sup>3</sup> | - |
| *COP1* (constitutive photomorphogenic ) | t007671 | AT4G11110 | 7:00 | 7:00 | 0 | 85 | Jan |
| *CRY1* (cryptochrome 1) | t007387 | AT4G08920 | 10:00 | 10:00 | 0 | 316 | Sep |
| *FT* (flowering locus T) | t021120 | AT4G20370 | 12:00 | 0:00 | 12 | 12 | Nov |
| *ELF4.2* (early flowering 4) | t027724 | AT1G72630 | 12:00 | 15:00 | 3 | 255 | Jul |
| *LHY* (late-elongated hypocotyl) | t004201 | AT2G46830 | 18:00 | 8:00 | 10 | *n.s.* | - |
| *GI* (gigantea) | t001582 | AT1G22770 | 18:00 | 20:00 | 2 | 97 | Apr |
| *ZTL* (Zeitlupe/F-box domain) | t005033 | AT2G18915 | 20:00 | 10:00 | 10 | *n.s.* | - |
| *TOC1* (timing of CAB expression) | t039766 | AT5G60100 | 20:00 | 19:00 | 1 | *n.s.* | - |
| *LUX* (myb-like transcription factor) | t012828 | AT5G59570 | 20:00 | 18:00 | 2 | *n.s.* | - |
| *ELF4.3* (early flowering 4) | t029722 | AT2G06255 | 20:00 | 20:00 | 0 | 249 | Jul |

<sup>1</sup> *Arabidopsis thaliana* phase estimates, as reported in Filichkin *et al.* Original values were published as hours from ZT (lights on), based on a 12 hour photoperiod. To make units comparable, we added 7 hours to *Arabidopsis* phase values to approximate the time of sunrise (in clock hours) in our study.
<sup>2</sup> Diurnal phase values are expressed clock hours, where sunrise occurred at 6:44 AM, and the photoperiod was 12:50 in length. Annual phase values are expressed in experiment days (days after 29-October).
<sup>3</sup> Profiles identified as not significantly rhythmic by JTK\_CYCLE are noted “*n.s.*”

From the 12,042 high-confidence diurnal transcripts, we were able to assign Mercator metabolic terms to 7,415 transcripts, and analyze these terms for overrepresentation or underrepresentation by categorizing transcriptional phases into bins representing four times of day: ‘sunrise’ (1,916 transcripts), ‘midday’ (1,333 transcripts), ‘sunset’ (2,685 transcripts), and ‘midnight’ (1,481 transcripts). Across daily bins, we found evidence for overrepresentation or underrepresentation in 15 Mercator metabolic pathways (Fig. 4A; supplementary file S3).

**Figure 4.**
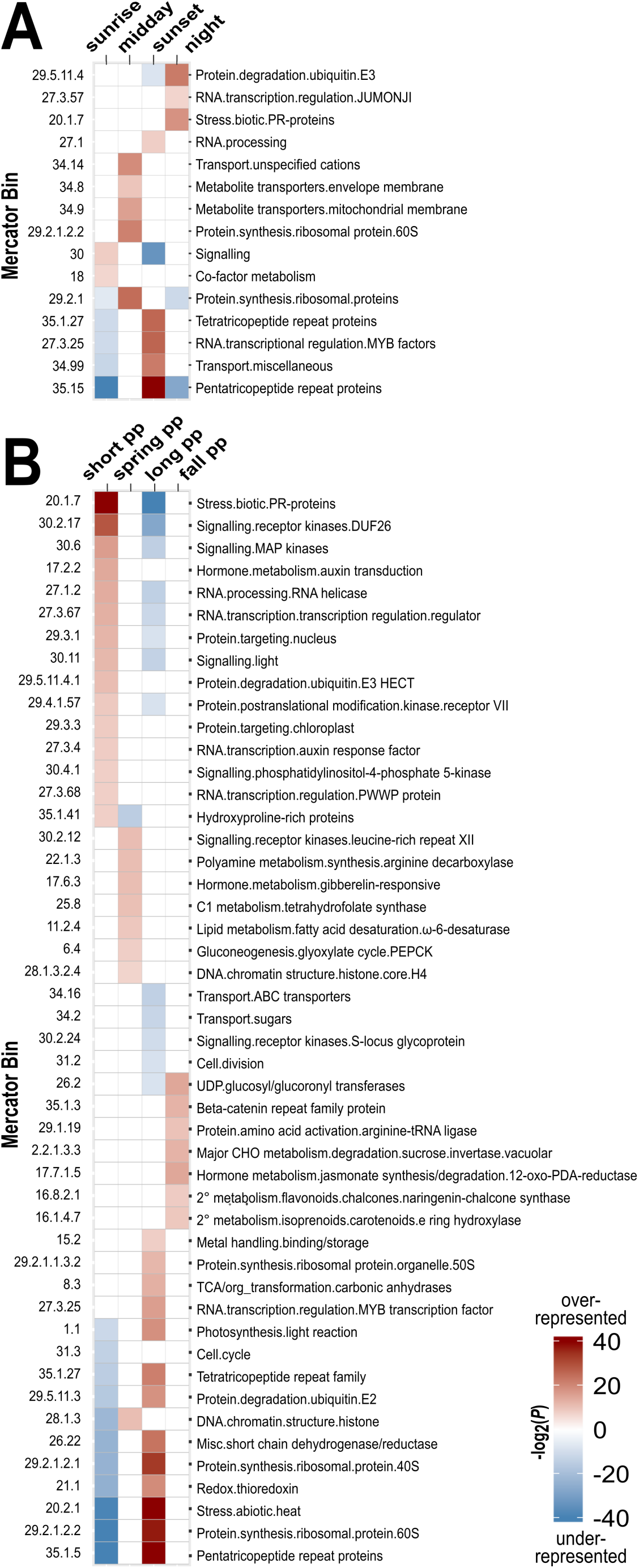
(A) Mercator metabolic categories showing evidence of significant enrichment (red) or depletion (blue) in the diurnal study. Transcripts were sorted by phase into one of four time bins; ‘sunrise’ (4 - 10am), ‘midday’ (10am – 4pm), ‘sunset’ (4 - 10pm), and ‘midnight’ (10pm – 4am). Shading is scaled by the −log2 value of the P-value, following an FDR correction of 0.05. (B) Metabolic pathways showing significant enrichment (red) or depletion (blue) in four photoperiod categories: short (5-Nov to 4-Feb), spring (5-Feb to 6-May), long (7-May to 5-Aug), and fall (6-Aug to 4-Nov). Shading is scaled by the −log2 value of the P-value, following an FDR correction of 0.05.

Overrepresented terms at sunrise include genes associated with light-responsive signaling (e.g., *phyB, cryl*, glutamate receptor-and cyclic nucleotide-gated ion channels proteins) and enzymes responsible for co-factor biosynthesis (e.g., biotin [holocarboxylase synthetase], thiamin [hydroxyethylthiazole kinase], CoA [phosphopantothenoylcysteine synthetase]). Overrepresented terms at midday include genes associated with protein synthesis (40S and 60S ribosomal proteins), and carbohydrate, nitrate, nucleotide, and small ion transporters. At sunset, overrepresented terms included diverse RNA modifying pathways, including pentatricopeptide repeat gene families (responsible for organelle RNA editing and processing), MYB transcription factors, and RNA processing genes (RNA pol I specific initiation factor *RRN3*; transducing/WD40 repeat proteins; mRNA decapping proteins; methyltransferases). Finally, overrepresented terms at midnight include genes associated with biotic stress (TIR-NBS-LRR proteins; LRR and NB-ARC proteins; *ADR1* -like proteins), *JUMONJI*-like histone demethylases known to play a role in the evening-phase of the *Arabidopsis* circadian clock, and protein degradation pathways based on ubiquitination/de-ubiquitination.

### Annual transcriptome variation in Douglas-fir tracks annual variation in photoperiod

Experiments to detect circannual variation included 5 individuals collected at 16 time points over 12 months, and the data were median filtered, normalized by DESeq, averaged by family, and then linearly interpolated to even sampling dates (Notes S1). After mapping reads from individual annual libraries to the pan-transcriptome reference, 36,145 transcripts met our threshold for analysis (Table 1). Following DESeq count normalization, JTK_CYCLE identified 24,688 transcripts showing significant circannual expression at a false-discovery rate of 5% (*q* ≤ 0.05). After imposing a more stringent false-discovery correction (*q* ≤ 0.01), the list contained 21,225 high-confidence circannual transcripts (Table 1; supplemental File S3).

The distribution of circannual expression phases (Fig. 2B) for high-coverage transcripts also shows a pronounced bimodal pattern, with the majority of transcripts reaching maximum expression in one of two seasons: (a) December through January, coinciding with winter dormancy, maximum freeze tolerance, reduced metabolic activity, and short photoperiods (day length ≤ 10.5 hours); or (b) June through July, coinciding with maximum shoot and radial growth, high metabolic activity, and long photoperiods (day length ≤ 14.5 hours). Example transcripts showing estimated phases for each month of the year are provided in Fig. 5, and details for these transcripts are provided in Notes 1. Included are transcripts homologous to genes known to play a role in winter adaptation and cold tolerance in *Arabidopsis* (Fig. 5A, inducer of CBP expression, *ICE1*), cold acclimation in potato (Fig 5B, stearoyl-ACP ∆9 desaturase gene; [22][60]), and proteins involved in winter photoprotection in conifers (Fig. 5C, early light inducible protein, *ELIP1;* [24]). Transcripts homologous to genes recently identified as showing evidence of convergent adaptation to cold climates in conifers [61] are also shown, including a mitochondrial transcription termination factor (Fig. 5F), a FMN-linked oxidoreductase (Fig. 5L), and a DNAJ-like heat shock protein (Fig. 5N). Transcripts homologous to genes related to QTLs strongly associated with drought in *Arabidopsis* [62] are also shown, including a cysteine proteinase (Fig. 5G), a pectin acetylesterase (Fig. 5H) and a chaperonin (Fig. 5K). As was observed with diurnal transcripts, 36.3% (*N* = 7,711) of the high-confidence annual cycling genes are “not assigned.unknown” in Mercator (Fig. 2B), and they show no homology to proteins or RNAs in GenBank. One example of a transcript with unknown function that is highly-expressed in spring is shown in Fig. 5D.

**Figure 5.**
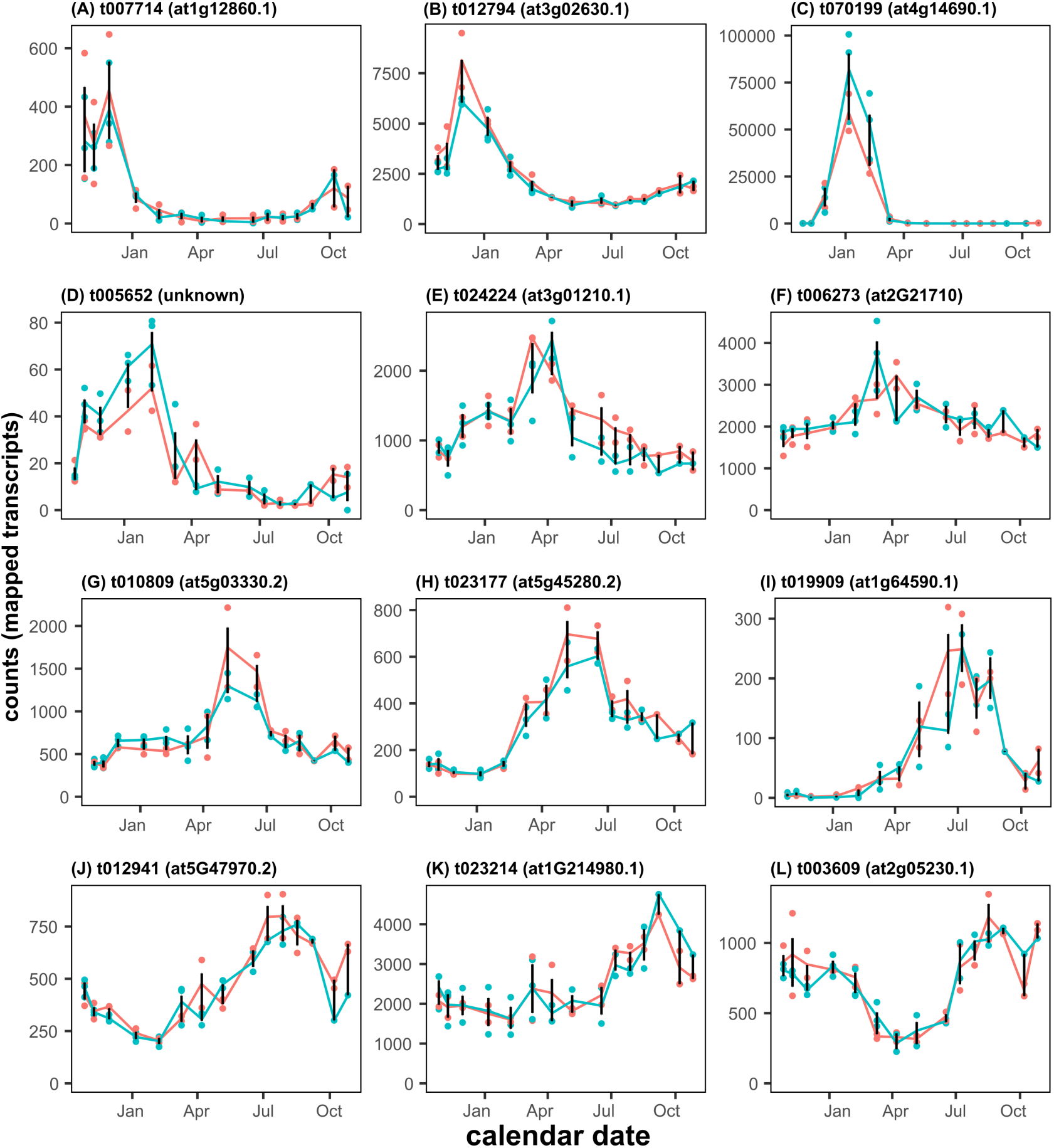
Example expression profiles for twelve transcripts exhibiting significant cyclic annual variation, ordered by monthly phase. Gene expression was estimated by RNA-seq from needle samples collected from two families of trees at *c*. three week intervals for one year. For each transcript, the Douglas-fir transcript name and best BLAST match to *Arabidopsis* (prefix ‘at’) are provided; genes lacking a BLAST match to the NCBI NR database are identified as ‘unknown’. Lines connect the mean gene expression (DESeq-normalized counts per transcript) for each family, and error bars represent the SD for all replicates.

We examined the relationship between the timing of maximum annual accumulation for transcription and four environmental inputs; mean photoperiod, biweekly cumulative precipitation, and bi-weekly mean temperature (maximum, minimum). Annual cyclic expression showed a strong significant association with photoperiod (*r*^2^ = 0.715, *F_24,2_* = 27.62, *P* ≤ 0.0001; Fig. 6A), and root mean square error was lower for a second-order polynomial fit than a linear fit. Annual cyclic expression also showed a significant but weaker association with bi-weekly cumulative precipitation (*r*^2^ = 0.368, *F*_24,2_ = 13.39, *P* ≤ 0.0011; Fig. 6B), and a lower root mean square error for linear versus a second-order polynomial fit. Annual cyclic expression showed non-significant associations with bi-weekly minimum and maximum temperatures (Fig. 6C-6D). Photoperiod and weather have been implicated to be major drivers in seasonal gene expression in different plant tissues [12][33]. Our results suggest that photoperiod is the dominant driver of the timing of cyclic gene expression maxima in Douglas-fir needles, but that precipitation and water availability is also positively associated with higher seasonal transcriptional activity.

**Figure 6.**
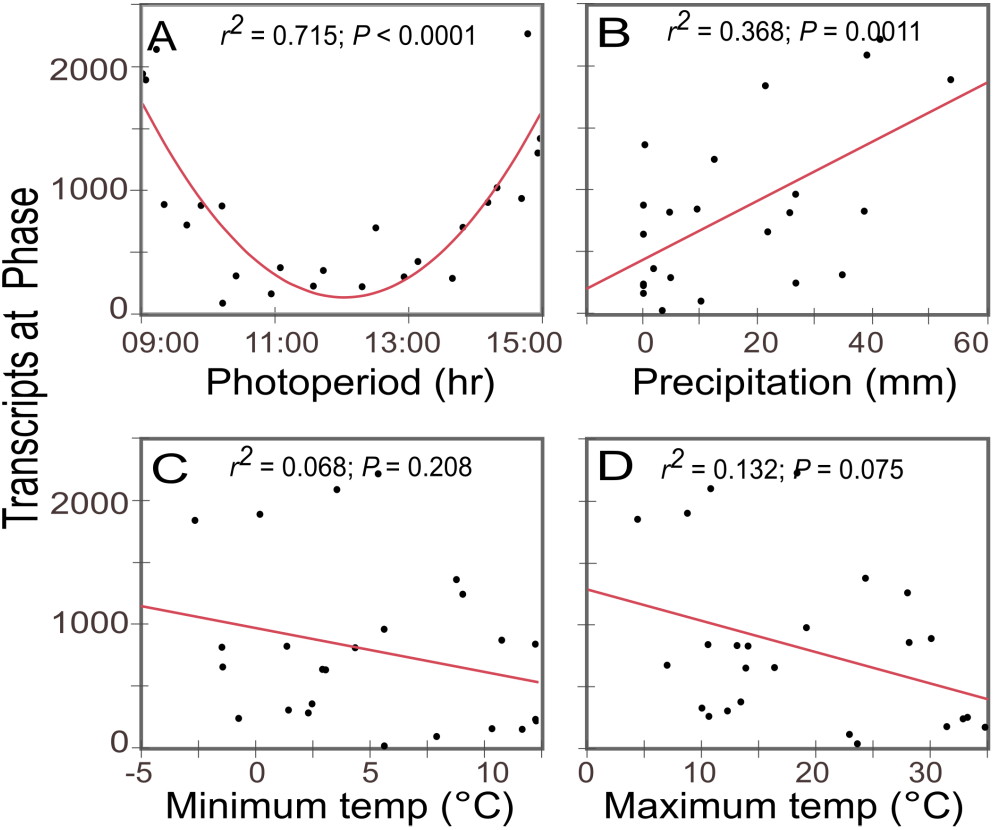
Regression results showing the relationship between the number of genes reaching expression maxima per two week period versus four environmental factors: (A) mean photoperiod; (B) mean solar radiation; (C) mean minimum temperature; and (D) two week cumulative precipitation. Regressions were performed using a polynomial fit (degree = 2) and best-fit lines are indicated in red.

Mercator terms were associated with 13,513 of the 21,225 high-confidence transcripts (excluding “not assigned.unknown”), and these were analyzed for evidence of over-representation or underrepresentation. Transcriptional phases were divided into seasonal bins by photoperiod (defined in Methods); bins included ‘short’ photoperiod (5,796 transcripts), ‘spring’ photoperiod (1,466 transcripts); ‘long’ photoperiod (5,073 transcripts); and ‘fall’ photoperiod (1,179 transcripts). Across seasonal bins, we found evidence for over-or underrepresentation in 48 Mercator metabolic pathways (Fig. 4B; supplemental file S4). Pathways showing an over-representation of phase values in short photoperiod days (winter) were related primarily to biotic stress, signaling, protein degradation and post-translational modification, and RNA transcription in diverse regulatory genes such as MAP kinases, RNA helicases, and proteins playing a role in hormone signaling and transduction (e.g., auxin; DUF26). Pathways showing over-representation of phases on long photoperiod days (summer) were related primarily to organelle regulation (pentatricopeptide and tetratricopeptide repeat protein families), photosynthesis-related metabolism (photosynthesis light reaction; carbonic anhydrase; metal binding/storage), ribosomal protein synthesis (nuclear and organellar), redox.thioredoxin, and stress from heat.

### Generalizing annual transcriptome expression variation to other conifers

If photoperiod is a dominant driver of rhythmic annual transcription in conifer needles, similar patterns of up- and down-regulation should be observable across other seasonal time-course studies of gene expression in conifers. To evaluate this hypothesis, we selected a subset of our annual data (17-Aug-2011 and 7-Oct-2011) to compare with a previously published microarray-based study of needle gene expression in Sitka spruce (*Picea sitchensis*) conducted at a similar time interval (30-Aug, 18-Oct) in British Columbia, Canada [33]. From these studies, we identified 11,582 transcripts that showed reciprocal best BLASTN matches between the Douglas-fir transcriptome and Sitka spruce clones used in microarray design (supplemental file S5). Out of this total, 760 transcripts showed evidence of significant seasonal change in Sitka Spruce (*q*≤0.05; two-fold change [33]), and cyclic behavior in Douglas-fir (*q*≤0.01, JTK-cycle). The rank order of fall:late summer gene expression ratios for these 760 transcripts was more preserved than would be expected if gene order was random (Kendall’s τ = 0.2901; z = 11.989; p < 2.2e-16), indicating that these experiments reveal similar patterns of fall season gene expression for a common set of genes, even though the comparisons are based on different evolutionary lineages (*Picea* vs. *Pseudotsuga)*, methodologies (microarray vs. RNA-seq), garden latitudes (42.3° N vs. 54° N), and sample dates. The general conservation of transcriptional patterns between these unrelated conifers highlights the utility of results from Douglas-fir as a tool for predicting annual expression trends and expression phase for homologous genes from other temperate zone conifers, such as spruce and pine [61].

## DISCUSSION

Perennial, evergreen needles are one of the key features that distinguish nearly 700 species of conifers from the tree model *Populus* and other deciduous angiosperm trees. Over a calendar year, persistent leaves undergo transcriptional and metabolic shifts that allow for high photosynthetic rates when conditions are favorable (even during winter), while providing conservative protection from the damaging effects of winter cold during dormancy, and drought during annual dry seasons [20]. By defining the complexity and contribution of gene expression over the complete growth-dormancy cycle, circannual studies like this provide a foundation for identifying associations between the synchrony of transcriptional change to seasonal changes and changes in phenology, and they offer a source of evidence for identifying genes/pathways that may contribute to adaptive responses in forest trees to climate change.

Our results show that the timing of maximum transcript accumulation in diurnally and circanually rhythmic transcripts from conifer needles is associated with the timing and quality of light at both temporal scales. At a daily scale, 12,042, or 29%, of expressed diurnal transcripts showed a significant diurnal rhythm, with two-thirds of these diurnal transcripts achieving maximum mRNA accumulation within +/− 2 hours of sunrise or sunset. At an annual scale, 21,225, or 58.7%, of expressed annual transcripts showed significant circannual rhythms, with nearly half of all expressed transcripts achieving maximum mRNA accumulation within +/− 20 days of the shortest or longest photoperiod. The influence of light as an entraining force of circadian rhythms is well-known in plants [1][8]; our analysis expands the list of known circadian genes in angiosperms to their lesser-studied sister lineage, the gymnosperms.

In contrast to daily cycles, the role of photoperiod on the timing of global transcript accumulation in perennial leaves is less well understood. Photoperiod is known to play a crucial role in the timing of the onset of dormancy, bud break and flowering in many plants [10][11][14]; our study shows that this ‘photoperiodic effect’ is a dominant pattern in the annual leaf transcriptome. Given the reliability of light as an entraining force for forecasting seasonal change, this is likely to be a common (if not universal) feature of temperate-zone conifer gene transcription. This hypothesis could be tested by evaluating cyclic transcriptome variation in trees grown in reciprocal gardens with fixed day length and contrasting temperature or precipitation regimes. In this type of comparison, genes showing light-dependent transcription should show a constant phase, irrespective of influences from the local climate. In contrast, genes responding to other exogenous cues like temperature and precipitation should show a shift in expression phase. These types of ‘needle-specific’ contrasts cannot be made with deciduous tree leaves, but equivalent comparisons could be made using other seasonally-responsive tissues (cambium; buds) from trees that are known to be responsive to photoperiod (e.g., *Populus* [9][12) and temperature (e.g., apple or pear [63]).

The entrainment of diurnal and circannual gene expression by light quality or day length in Douglas-fir is intuitive, but the dramatic accumulation of transcripts in winter presents a paradox; *why does transcript accumulation peak for such a high proportion of the cyclic transcriptome during dormancy, when metabolic activity is reduced and growth is arrested?* In western Oregon, Douglas-fir remains photosynthetically active during the winter [20], but it undergoes physiological changes that result in maximum cold hardiness between November to early December [23]; this timing coincides with the increase in genes achieving maximum transcript accumulation (e.g., Figure 2B). Transcription for many genes and pathways are known to increase under short days and cold temperatures, including those that show associations to winter cold-acclimation responses [24] [22]. Direct comparisons to leaf transcriptomes cannot be made with winter-deciduous angiosperms, but the list of Douglas-fir cyclic transcripts reaching maximum transcription in winter includes genes that have been implicated in cold acclimation of *Populus* buds (e.g., early light-inducible proteins (ELIPs), C-repeat binding factors, fatty acid desaturases, major carbohydrate enzymes, LEA-like proteins, and heat shock proteins [17]. Short day-induced transcriptional upregulation in needles appears extensive, and further effort is required to determine whether this massive increase in transcript accumulation is a response to the proximal demands of growth cessation, dormancy, cold hardiness, or more distant processes involved in breaking dormancy.

Our study highlights a related paradox, which is that the physiologically complex events occurring during the resumption of spring growth (increased photosynthetic rate; onset of radial and shoot growth; flowering) occur without a large, coordinated transcriptional response from cyclic genes in needles. This suggests that the transcriptional activity required to end dormancy and resume spring growth involves a comparatively small number of cyclic genes and pathways, that the activity is partitioned to other tissues and tissue-specific cyclic genes, or that the coordinated response occurs far in advance of the actual event of spring growth. The veracity of these alternatives could be readily evaluated using comparative time-series gene expression (or proteomic) experiments using different tissues from individual plants over the dormancy-growth transition.

The daily and circannual transcriptional patterns for Douglas-fir are available for examination [64], and these will be useful for predicting the timing of transcription in genomically-complex gymnosperm models such as spruce [65][66] and pine [48]. The combination of large genome size (~20 Gbp), high transcriptional complexity [67][68], genic redundancy, and divergence from angiosperm models has made it difficult to infer gene function in conifers based on homology alone, although genetic-environmental and genetic-phenotypic associations are being investigated in many conifer species [61] [69]. Conifers lack tractable models for reverse genetic manipulation, so context-specific knowledge offered by diurnal, seasonal, and circannual gene expression studies can provide additional clues to the timing of transcript accumulation, and the seasonal context by which genes and pathways are up- and downregulated. Accurate estimates of phase, and changes in phase under different experimental conditions, will provide a means to characterize and define regulatory relationships in pathways critical to a variety of functions. A clearer understanding of these complex circannual responses will emerge as the developing Douglas-fir genome [70] and other conifer genomes are integrated with daily, seasonal, circannual, and tissue-specific transcriptomic studies. This knowledge will set the stage for understanding how conifer needles have responded to climate changes during their 300+ million year independent history from flowering plants, and it may offer novel avenues for predicting adaptive responses under different future climate alternatives.

## DECLARATIONS

### Availability of data and material

The sequences, reference transcriptome and mapped count data are available under NCBI BioProject Umbrella **PRJNA362352;** this includes diurnal sequences and counts (**PRJNA263611**), annual diurnal sequences and counts (**PRJNA243096**), and reference transcriptome (**PRJNA356432**). Mercator and GO annotations are available from the TreeGenes website (http://treegenesdb.org/ftp/Transcriptome_Data/transcriptome/Psme/Cyclic_Transcriptome_Assembly_v2.04_%5bFS%5d/), and data sets analyzed for this published article are available as supplementary files.

### Authors' contributions

R.C.C., PCD., J.B.S. and D.R.D. jointly designed the study; R.C.C., P.C.D. and J.B.S. performed experiments and collected data; R.C.C., P.C.D. and S.H. analyzed data; S.J., J.L.W. and D.B.N. provided draft transcriptome and genome assemblies, and provided transcript annotations; R.C.C., P.C.D., S.J., B.S.C. and D.R.D. wrote the manuscript, and J.L.W. and D.B.N. gave conceptual advice on the manuscript. All authors edited and approved the final version of the manuscript.

## Acknowledgements

We thank Peter Gould (Washington Department of Natural Resources), Constance Harrington (US Forest Service), and Brian Knaus (USDA Agricultural Research Service) for input on garden design, sampling strategy, and general advice. Tara Jennings, Brian Knaus, Danielle Marca and Chris Poklemba assisted with tissue collections, and Katie Alderman, Tara Jennings and Laura Mealy assisted with RNA extraction and library construction. Mark Dasenko, Matthew Peterson and Chris Sullivan (Oregon State University) performed Illumina sequencing, and Hans Vasquez-Gross (University of California-Davis) assisted with transcript annotation. We thank Engin Sunger (University of Minnesota-Morris) and the Michael Hughes laboratory (University of Missouri - St. Louis) for providing advice on using JTK-cycle with irregularly-spaced samples, and Katherine Hayden, Glenn Howe, Hardeep Rai, and Jessica Wright for input and advice at early stages of this project.

## The following Supporting Information is provided for this article

### Notes S1 Additional File 1

Addition-al_File_1.rtf. Additional information on needle sampling methods, original source locations for trees, the location of common gardens, sampling intervals used for collections, individual tree sequencing library summaries, and individual tree transcriptome summaries. The file can be opened in Microsoft Word or any word processing program that accepts rich text format (RTF).

### File S1 Additional File 2

Additional_File_2.tsv. Daily environmental data associated with experiment. This file includes date (mm/dd/yyyy), experiment day, Julian day, experiment week, experiment month, minimum air temperature (°C), maximum air temperature (°C), mean daily air temperature (° C), daily precipitation (mm), day length (h:m:s), and the sum of solar intensity (megajoules/m^2^). The file can be opened in Microsoft EXCEL or any program that accepts text as tab-separated values (TSV).

### File S2 Additional File 3

Additional_File_3.tsv. Summary of JTK-CY CLE inferred cycle properties for diurnal and annual cyclic transcripts. The file can be opened in Microsoft EXCEL or any program that accepts text as tab-separated values (TSV).

### File S3 Additional File 4

Addition-al_File_4.xlsx. Summary of enriched and depleted Mercator functional category terms for the diurnal study. The file can be opened in Microsoft EXCEL (XLS).

### File S4 Additional File 5

Additional_File_5. xlsx. Summary of enriched and depleted Mercator functional category terms for the annual study. The file can be opened in Microsoft EXCEL (XLS).

### File S5 Additional File 6

Additional_File_6. xlsx. Summary of transcripts showing best reciprocal BLAST match between Douglas-fir and Sitka spruce clones used in Holliday et al., 2008, and data used for comparing ranked lists for Kendall’s Tau test. The file can be opened in Microsoft EXCEL (XLS).

## REFERENCES

1. Bell-Pedersen D, Cassone VM, Earnest DJ, Golden SS, Hardin PE, Thomas TL, et al. Circadian rhythms from multiple oscillators: lessons from diverse organisms. Nat. Rev. Genet. 2005;6:544–56.

2. Dodd AN. Plant circadian clocks increase photosynthesis, growth, survival, and competitive advantage. Science. 2005;309:630–3.

3. McClung CR. Plant circadian rhythms. Plant Cell. 2006;18:792–803.

4. Pruneda-Paz JL, Kay SA. An expanding universe of circadian networks in higher plants. Trends Plant Sci. 2010;15:259–65.

5. Hughes ME, Grant GR, Paquin C, Qian J, Nitabach MN. Deep sequencing the circadian and diurnal transcriptome of *Drosophila* brain. Genome Res. 2012;22:1266–81.

6. Dopico XC, Evangelou M, Ferreira RC, Guo H, Pekalski ML, Smyth DJ, et al. Widespread seasonal gene expression reveals annual differences in human immunity and physiology. Nat. Commun. 2015;6:7000.

7. Filichkin SA, Breton G, Priest HD, Dharmawardhana P, Jaiswal P, Fox SE, et al. Global profiling of rice and poplar transcriptomes highlights key conserved circadian-controlled pathways and cis-regulatory modules. PLoS ONE. 2011;6:e16907.

8. Nagel DH, Doherty CJ, Pruneda-Paz JL, Schmitz RJ, Ecker JR, Kay SA. Genome-wide identification of CCA1 targets uncovers an expanded clock network in *Arabidopsis*. Proc. Natl. Acad. Sci. 2015;112:E4802–E4810.

9. Ingvarsson PK, García MV, Hall D, Luquez V, Jansson S. Clinal variation in phyB2, a candidate gene for day-length-induced growth cessation and bud set, across a latitudinal gradient in European Aspen *(Populus tremula)*. Genetics. 2006;172:1845–53.

10. Horvath DP, Anderson JV, Chao WS, Foley ME. Knowing when to grow: signals regulating bud dormancy. Trends Plant Sci. 2003;8:534–40.

11. Böhlenius H, Huang T, Charbonnel-Campaa L, Brunner AM, Jansson S, Strauss SH, et al. CO/FT regulatory module controls timing of flowering and seasonal growth cessation in trees. Science. 2006;312:1040–3.

12. Yordanov YS, Ma C, Strauss SH, Busov VB. EARLY BUD-BREAK 1 (EBB1) is a regulator of release from seasonal dormancy in poplar trees. Proc. Natl. Acad. Sci. 2014;111:10001–6.

13. Peñuelas J, Filella I. Responses to a warming world. Science. 2001;294:793–5.

14. Lagercrantz U. At the end of the day: a common molecular mechanism for photoperiod responses in plants? J. Exp. Bot. 2009;60:2501–15.

15. Olsen JE. Light and temperature sensing and signaling in induction of bud dormancy in woody plants. Plant Mol. Biol. 2010;73:37–47.

16. Yu H, Luedeling E, Xu J. Winter and spring warming result in delayed spring phenology on the Tibetan Plateau. Proc. Natl. Acad. Sci. 2010;107:22151–6.

17. Rohde A, Ruttink T, Hostyn V, Sterck L, Van Driessche K, Boerjan W. Gene expression during the induction, maintenance, and release of dormancy in apical buds of poplar. J. Exp. Bot. 2007;58:4047–60.

18. Harrington CA, Gould PJ, St. Clair JB. Modeling the effects of winter environment on dormancy release of Douglas-fir. For. Ecol. Manag. 2010;259:798–808.

19. Ruttink T, Arend M, Morreel K, Storme V, Rombauts S, Fromm J, et al. A molecular timetable for apical bud formation and dormancy induction in poplar. Plant Cell. 2007;19:2370–90.

20. Waring RH, Franklin JF. Evergreen coniferous forests of the *Pacific Northwest*. Science. 1979;204:1380–6.

21. Demmig-Adams B, Adams WW. Photoprotection in an ecological context: the remarkable complexity of thermal energy dissipation. New Phytol. 2006;172:11–21.

22. Mayr S, Schmid P, Laur J, Rosner S, Charra-Vaskou K, Damon B, et al. Uptake of water via branches helps timberline conifers refill embolized xylem in late winter. PLANT Physiol. 2014;164:1731–40.

23. Aitken SN, Adams WT. Spring cold hardiness under strong genetic control in Oregon populations of *Pseudotsuga menziesii* var. menziesii. Can. J. For. Res. 1997;27:1773–80.

24. Zarter CR, Adams WW, Ebbert V, Cuthbertson DJ, Adamska I, Demmig-Adams B. Winter down-regulation of intrinsic photosynthetic capacity coupled with up-regulation of *elip-like* proteins and persistent energy dissipation in a subalpine forest. New Phytol. 2006;172:272–82.

25. Gernandt DS, Magallón S, López GG, Flores OZ, Willyard A, Liston A. Use of simultaneous analyses to guide fossil-based calibrations of Pinaceae phylogeny. Int. J. Plant Sci. 2008;169:1086–99.

26. Chen P-Y, Welsh C, Hamann A. Geographic variation in growth response of Douglas-fir to interannual climate variability and projected climate change. Glob. Change Biol. 2010;16:3374–85.

27. Campbell RK, Sorensen FC. Cold-acclimation in seedling Douglas-Fir related to phenology and provenance. Ecology. 1973;54:1148–51.

28. St. Clair JB, Mandel NL, Vance-Borland KW. Genecology of Douglas-fir in western Oregon and Washington. Ann. Bot. 2005;96:1199–214.

29. Anekonda TS, Adams WT, Aitken SN, Neale DB, Jermstad KD, Wheeler NC. Genetics of cold hardiness in a cloned full-sib family of coastal Douglas-fir. Can. J. For. Res. 2000;30:837–840.

30. Bansal S, St. Clair JB, Harrington CA, Gould PJ. Impact of climate change on cold hardiness of Douglas-fir (*Pseudotsuga menziesii*): environmental and genetic considerations. Glob. Change Biol. 2015;21:3814–26.

31. Hughes ME, Hogenesch JB, Kornacker K. JTK_CYCLE: an efficient non-parametric algorithm for detecting rhythmic components in genome-scale datasets. J. Biol. Rhythms. 2010;25:372–80.

32. Bar-Joseph Z, Gitter A, Simon I. Studying and modelling dynamic biological processes using time-series gene expression data. Nat. Rev. Genet. 2012;13:552–64.

33. Holliday JA, Ralph SG, White R, Bohlmann J, Aitken SN. Global monitoring of autumn gene expression within and among phenotypically divergent populations of Sitka spruce *(Picea sitchensis)*. New Phytol. 2008;178:103–22.

34. Gould PJ, Harrington CA, Clair JBS. Growth phenology of coast Douglas-fir seed sources planted in diverse environments. Tree Physiol. 2012;32:1482–96.

35. Tai H, Pelletier C, Beardmore T. Total RNA isolation from *Picea mariana* dry seed. Plant Mol. Biol. Report. 2004;22:93a–93e.

36. Parkhomchuk D, Borodina T, Amstislavskiy V, Banaru M, Hallen L, Krobitsch S, et al. Transcriptome analysis by strand-specific sequencing of complementary DNA. Nucleic Acids Res. 2009;37:e123.

37. Bolger A, Lohse M, Usadel B. Trimmomatic: A flexible trimmer for Illumina sequence data. Bioinformatics. 2014;30:2114–20.

38. Grabherr MG, Haas BJ, Yassour M, Levin JZ, Thompson DA, Amit I, et al. Full-length transcriptome assembly from RNA-Seq data without a reference genome. Nat. Biotechnol. 2011;29:644–52.

39. Edgar RC. Search and clustering orders of magnitude faster than BLAST. Bioinformatics. 2010;26:2460–1.

40. TreeGenes: A Forest Tree Genome Database [Internet]. TreeGenes For. Tree Genome Database. [cited 2016 Nov 15]. Available from: https://dendrome.ucdavis.edu/treegenes/species/index.php?letter=Pseudotsuga&#results

41. Altschul SF, Gish W, Miller W, Myers EW, Lipman DJ. Basic local alignment search tool. J. Mol. Biol. 1990;215:403–10.

42. Lohse, M, Nagel A, Herter T, May P, Schroda M, Zrenner R, et al. Mercator: a fast and simple web server for genome scale functional annotation of plant sequence data. Plant Cell Environ. 2014;37:1250–8.

43. Klie S, Nikoloski Z. The choice between MapMan and Gene Ontology for automated gene function prediction in plant science. Front. Genet. [Internet]. 2012 [cited 2016 Aug 16];3. Available from: http://journal.frontiersin.org/article/10.3389/fgene.2012.00115/abstract

44. Mercator pipeline for automated sequence annotation [Internet]. Mercat. Pipeline Autom. Seq. Annot. [cited 2016 Nov 15]. Available from: http://mapman.gabipd.org/web/guest/app/mercator

45. Harris R. Improved pairwise alignment of genomic DNA. PhD thesis, The Pennsylvania State University, State College, PA, USA. ISBN 978-0-549-43170-1 [Inter.net]. [State College, PA]: The Pennsylvania State University; 2007 [cited 2016 Mar 11]. Available from: http://www.bx.psu.edu/~rsharris/rsharris_phd_thesis_2007.pdf

46. Index of /ftp/Genome_Data/genome/pinerefseq/Pita/mito [Internet]. Pinus Taeda Mitochondrial Scaffold Seq. Version 5112015. [cited 2016 Nov 15]. Available from: http://dendrome.ucdavis.edu/ftp/Genome_Data/genome/pinerefseq/Pita/mito/

47. Kent WJ. BLAT — The BLAST-like alignment tool. Genome Res. 2002;12:656–64.

48. Neale DB, Wegrzyn JL, Stevens KA, Zimin AV, Puiu D, Crepeau MW, et al. Decoding the massive genome of loblolly pine using haploid DNA and novel assembly strategies. Genome Biol. 2014;15:R59.

49. Index of /ftp/Genome_Data/genome/pinerefseq/Psme [Internet]. 2016 [cited 2016 Nov 15]. Available from: http://treegenesdb.org/ftp/Transcriptome_Data/transcriptome/Psme/Cyclic_Transcriptome_Assembly_v2.04_%5bFS%5d/

50. Langmead B, Salzberg SL. Fast gapped-read alignment with Bowtie 2. Nat. Methods. 2012;9:357–9.

51. Anders S, Huber W. Differential expression analysis for sequence count data. Genome Biol. 2010;11:R106.

52. Bullard JH, Purdom E, Hansen KD, Dudoit S. Evaluation of statistical methods for normalization and differential expression in mRNA-Seq experiments. BMC Bioinformatics. 2010;11:94.

53. Dillies M-A, Rau A, Aubert J, Hennequet-Antier C, Jeanmougin M, Servant N, et al. A comprehensive evaluation of normalization methods for Illumina high-throughput RNA sequencing data analysis. Brief. Bioinform. 2013;14:671–83.

54. Benjamini Y, Hochberg Y. Controlling the false discovery rate: a practical and powerful approach to multiple testing. J. R. Stat. Soc. Ser. B. 1995;57:289–300.

55. Giorgi F. MEFISTO: MapMan Enrichment via FISher’s Test for Ontology. [WWW document] URL http://www.usadellab.org/cms/index.php?page=corto. [accessed 12 March 2016]. [Internet]. 2012. Available from: http://www.usadellab.org/cms/index.php?page=corto

56. Rivals I, Personnaz L, Taing L, Potier M-C. Enrichment or depletion of a GO category within a class of genes: which test? Bioinformatics. 2007;23:401–7.

57. Coate JE, Doyle JJ. Quantifying whole transcriptome size, a prerequisite for understanding transcriptome evolution across species: an example from a plant allopolyploid. Genome Biol. Evol. 2010;2:534–46.

58. Lovén J, Orlando DA, Sigova AA, Lin CY, Rahl PB, Burge CB, et al. Revisiting global gene expression analysis. Cell. 2012;151:476–82.

59. Gyllenstrand N, Clapham D, Källman T, Lagercrantz U. A Norway Spruce FLOWERING LOCUS T homolog is implicated in control of growth rhythm in conifers. Plant Physiol. 2007;144:248–57.

60. Vega SE, del Rio AH, Bamberg JB, Palta JP. Evidence for the up-regulation of stearoyl-ACP (Δ9) desaturase gene expression during cold acclimation. Am. J. Potato Res. 2004;81:125–35.

61. Yeaman S, Hodgins KA, Lotterhos KE, Suren H, Nadeau S, Degner JC, et al. Convergent local adaptation to climate in distantly related conifers. Science. 2016;353:1431–3.

62. Bac-Molenaar JA, Granier C, Keurentjes JJB, Vreugdenhil D. Genome-wide association mapping of time-dependent growth responses to moderate drought stress in *Arabidopsis*. Plant Cell Environ. 2016;39:88–102.

63. Heide OM, Prestrud AK. Low temperature, but not photoperiod, controls growth cessation and dormancy induction and release in apple and pear. Tree Physiol. 2005;25:109–14.

64. Dolan P. Douglas-fir Transcriptome Data [Internet].[cited 2016 Nov 15]. Available from: http://146.57.34.125:3838/

65. Birol I, Raymond A, Jackman SD, Pleasance S, Coope R, Taylor GA, et al. Assembling the 20 Gb white spruce (*Picea glauca*) genome from whole-genome shotgun sequencing data. Bioinformatics. 2013;29:1492–7.

66. Nystedt B, Street NR, Wetterbom A, Zuccolo A, Lin Y-C, Scofield DG, et al. The Norway spruce genome sequence and conifer genome evolution. Nature. 2013;497:579–84.

67. Howe GT, Yu J, Knaus B, Cronn R, Kolpak S, Dolan P, et al. A SNP resource for Douglas-fir: de novo transcriptome assembly and SNP detection and validation. BMC Genomics. 2013;14:137.

68. Wegrzyn JL, Liechty JD, Stevens KA, Wu L-S, Loopstra CA, Vasquez-Gross HA, et al. Unique features of the Loblolly Pine *(Pinus taeda* L.) megagenome revealed through sequence annotation. Genetics. 2014;196:891–909.

69. Eckert AJ, Bower AD, Wegrzyn JL, Pande B, Jermstad KD, Krutovsky KV, et al. Association genetics of coastal Douglas Fir (*Pseudotsuga menziesii* var. *menziesii*, Pinaceae). I. Cold-hardiness related traits. Genetics. 2009;182:1289–302.

70. PineRefSeq: Pine Reference Sequences [Internet]. [cited 2016 Nov 15]. Available from: http://pinegenome.org/pinerefseq/

